# Latitudinal cline in hypoxia tolerance does not result in correlated acid tolerance in *Tigriopus californicus*

**DOI:** 10.1101/2021.04.10.439290

**Authors:** Aimee Deconinck, Christopher Willett

## Abstract

Intertidal organisms must tolerate a wide range of environmental parameters each day which may result in tolerance to multiple stressors correlating. The intertidal copepod *Tigriopus californicus* experiences diurnal variation in dissolved oxygen levels and pH as the opposing processes of photosynthesis and cellular respiration lead to coordinated highs during the day and lows at night. While environmental parameters with overlapping spatial gradients frequently result in correlated traits, less attention has been given to exploring temporally correlated stressors. We investigated whether hypoxia tolerance correlated with acid tolerance by testing the hypoxia and low pH stress tolerance of 6 genetically differentiated populations of *T. californicus*. We checked for similarities in tolerance for the two stressors by latitude, sex, size, and time since collection as predictors. We found that although hypoxia tolerance correlated with latitude, acid tolerance did not, and no predictor was significant for both stressors. We concluded that temporally coordinated exposure to low pH and low oxygen did not result in populations developing equivalent tolerance for both.

## Introduction

Physiological tolerance is essential in highly variable environments, and few marine inhabitants experience as much routine environmental change as residents of the rocky intertidal habitat. High intertidal rock pools experience daily and seasonal changes in temperature, dissolved oxygen (DO), and pH rarely seen in other marine systems (Morris & Taylor, 1983; Truchot & Duhamel-Jouve, 1980; Wolfe, Worjanyn, & Byrne, 2013). In surface aquatic systems, DO and pH covary temporally as the opposing processes of photosynthesis and aerobic respiration drive the exchange of oxygen and carbon dioxide gases. During the day, photosynthetic organisms use dissolved carbon dioxide gas for photosynthesis and release oxygen gas, raising DO levels. The use of dissolved carbon dioxide also shifts the equilibrium of the CO_2_-bicarbonate-carbonate system, increasing the pH (Hutchinson, 1957). At night, respiration consumes oxygen and releases carbon dioxide, reversing the pattern to the point where waters may become hypoxic (DO ≤2.0 mg O_2_/L or 30% saturation) and slightly acidic (pH <7) (Morris & Taylor, 1983). When species respond to multiple environmental gradients in the absence of genetic constraints, phenotypes are predicted to correlate as well (Duputié, Massol, Chuine, Kirkpatrick, & Ronce, 2012; Futuyma & Moreno, 1988; Hoffmann & Parsons, 1991). Thus, we might expect tolerance to decreased oxygen levels and decreased pH to correlate as well.

When phenotypic differences arise due to selection in different environments and are not the result of plasticity or drift, local adaptation may be involved (De Villemereuil, Gaggiotti, Mouterde, & Till-Bottraud, 2016; Duputié et al., 2012; Nicolaus & Edelaar, 2018). *Tigriopus californicus* inhabit high intertidal splash pools from the central Baja Peninsula in Mexico to southern Alaska in the US. which includes a wide range of physiochemical properties. These geographic differences in habitat combined with low dispersal have resulted in populations with highly divergent genomes (Edmands, 2001; Burton *et al*., 2007; Barreto *et al*., 2018, Table S1). Common garden experiments have demonstrated local adaptation to thermal tolerance and some measures of salinity tolerance align with latitude, as southern populations are more tolerant of heat shock (M. W. Kelly, Sanford, & Grosberg, 2012; Pereira, Sasaki, & Burton, 2017; Willett, 2010) and northern populations, which receive more rain, are generally more tolerant of extended periods at low salinity (Leong, Sun, & Edmands, 2018, although see also J. Lee, Phillips, Lobo, & Willett, 2021).

Although we are unaware of any studies specifically exploring latitudinal clines of hypoxia and acid adaptation, multiple studies suggest that temperature, which does correspond with latitude, influences hypoxia and acid tolerance. *Tigriopus brevicornis* from Scotland consumed less oxygen at low temperatures while experiencing hypoxia, but at the highest temperature the level of oxygen needed to maintain oxygen regulation (P_c_) was significantly elevated (McAllen, Taylor, & Davenport, 1999). Although *T. californicus* from Washington survived high temperature stress approximately the same regardless of DO levels, the lowest DO level tested was 22% of saturation for 5 hours (Dinh, Cuevas-Sanchez, Buhl, Moeser, & Dowd, 2020). In another study, *T. californicus* copepods survived at least 24 hours at anoxia (Graham & Barreto, 2019), which may indicate that the DO levels tested by Dinh *et al.* were not stressful. Several copepod species exhibit signs of sublethal stress to ocean acidification when it is paired with higher temperatures (Wang, Jeong, Lee, & Lee, 2018). Taken with the observation that warmer water dissolves less oxygen than colder water, these studies suggest that elevated temperatures can increase hypoxia and acid stress. When gene flow is limited, as seen between *T. californicus* populations, local adaptation should be favored over plasticity when there is spatial structure to patterns of environmental variability (Sultan & Spencer, 2002). Thus, we expected that southern populations of *T. californicus* would experience more intense hypoxia and low pH stress and would have higher tolerance to the stressors than northern populations.

In addition to latitudinal clines in environmental parameters, stress tolerance may also be associated with sex-specific adaptation, laboratory adaptation and life history traits. Sex-specific variation in environmental stress tolerance is well documented, and, in *T. californicus*, males are often the more sensitive sex (Foley et al., 2019). However, models comparing directional to cyclic environmental change predict that the difference between sexes should decrease with rapidly changing environments (Connallon & Hall, 2016). Adaptation to laboratory conditions are also well documented, as populations respond to different selective pressures than they experienced in the field (Hoffmann, Hallas, Sinclair, & Partridge, 2001; Hoffmann & Ross, 2018; Kingsolver & Nagle, 2007). In contrast to the unidirectional relationship between stress and sex-specific or laboratory adaptation, body size differences may be both a response to environmental stress (Atkinson, 1994; de Oliveira Sodré & Bozelli, 2019; Rowiński & Rogell, 2017) as well as a predictor of thermal tolerance (Nielsen & Papaj, 2015; Rubalcaba & Olalla-Tárraga, 2020), although its impact on other forms of stress are less clear (Nilsson & Östlund-Nilsson, 2008; Schmitz & Harrison, 2004; Spicer & Morley, 2019). Since multiple predictors may influence stress tolerance, this paper aims to determine which variables are most important for predicting hypoxia and acid tolerance in genetically divergent populations of *T. californicus*.

We tested adult copepods for either hypoxia tolerance or acid tolerance to determine if diurnal correlation of the stressors resulted in phenotype correlation. We expected (1) the frequency of these stressors co-occurring temporally would result in population-specific tolerance to both stressors, (2) hypoxia and acid tolerance would inversely correlate with latitude, and (3) sex, body size, and time since collection would have significant effects on tolerance.

## Methods and Materials

Populations of *Tigriopus californicus* were collected between September 2004 and April 2018 from 6 rocky intertidal locations along the eastern Pacific coastline: Friday Harbor Laboratories, WA; Bodega Marine Lab, CA; Santa Cruz, CA; San Simeon, CA; Abalone Cove, CA; and San Diego, CA (Table 1 and Table S1). After maintaining populations under the same environmental conditions for at least 6 generations (see supplement for rearing details), individual copepods were used only once in a single-stressor trial for the hypoxia assay, low pH assay or control run.

**Table 1.** Collection locations, dates, and sample sizes.

| Population | Abbreviation | Latitude,<br>Longitude | Date of<br>Collection<br>(MM/DD/YY) | Sample Size:<br>Hypoxia (sex)<br>Acid (sex) |
| --- | --- | --- | --- | --- |
| Friday Harbor<br>Laboratories | FHL | 48.449974,<br>-122.963021 | 4/3/18 | 114 (57 f, 57 m)<br>80 (40 f, 40 m) |
| Bodega Marine Lab | BB | 38.305133,<br>-123.065556 | 7/9/13 | 119 (60 f, 59 m)<br>79 (40 f, 39 m) |
| Santa Cruz | SCN | 36.949617,<br>-122.046940 | 11/21/17 | 120 (60 f, 60 m)<br>80 (40 f, 40 m) |
| San Simeon | SS | 35.581633,<br>-121.121111 | 9/27/04 | 114 (55 f, 59 m)<br>79 (40 f, 39 m) |
| Abalone Cove | AB | 33.740733,<br>-118.377778 | 5/24/11 | 116 (60 f, 56 m)<br>77 (38 f, 39 m) |
| San Diego | SD | 32.745646,<br>-117.255036 | 11/22/17 | 113 (55 f, 58 m)<br>77 (39 f, 38 m) |

### Hypoxia and low pH assays

Both the hypoxia assay and the low pH assay were conducted in a custom-built glove box (Figure S1). During each assay, up to five copepods were grouped by population and sex and placed in a single arena made of a plastic cylinder with a fine mesh bottom. Eight arenas were placed in a water bath that was maintained at 20°C (±2°C) and enclosed by the glove box. Populations and sexes were assigned randomly to one of the eight arenas during each trial to minimize placement effects. In all assays, animals were not fed during the gas adjustment period or the experiment.

For the hypoxia assay, nitrogen gas was bubbled continuously to displace dissolved oxygen and maintain a low oxygen environment followed by a reoxygenation period before measuring survival. The hypoxic period was started when the DO level was ≤0.1 mg/L which took 1.98±0.60 hours of bubbling nitrogen gas to reach 0.05 mg/L. Although hypoxia is defined as ≤2 mg/L, exploratory experiments indicated that *T. californicus* were highly tolerant of DO levels greater than 0.1 mg/L (unpublished data). Hypoxia was maintained for 20 hours, and dissolved oxygen was measured using an Orion Star A213 Dissolved Oxygen (DO) Benchtop Meter (Thermo Scientific) throughout the experiment. After the hypoxic period, atmospheric air was pumped into the box for 10 hours using an aquarium air pump, then the number of surviving individuals were counted by gently prodding individuals with a pipette tip to elicit an evasion response (swimming away).

For the low pH assay, carbon dioxide was bubbled into the water bath until the pH measured 4.8 then the water bath was allowed to equilibrate with the atmosphere for 24 hours before measuring survival. Preliminary work showed this elicited a knockdown response in the majority of samples (unpublished data). When the pH reached 4.8 which took less than 10 minutes, the gas supply was turned off, and the water bath was allowed to return to normal pH over time (Figure S2). The number of swimming individuals, temperature, pH and DO were sampled every 20 minutes for the first 6 hours, then again at 12 hours and 24 hours using the same protocol described above. Dissolved oxygen levels did not drop below 1.85 mg/L during any trial.

Control trials for both assays were set up with the same materials and conducted at the same time as the hypoxia and low pH assays except atmospheric air was added using an aquarium air pump rather than nitrogen or carbon dioxide gas. Due to the limitation of equipment, DO and pH for control trials were only sampled at the beginning and end of an assay, although values remained consistent (pH = 7.81±0.30; DO = 8.22±0.26 mg/L). When possible, siblings from the same brood were split between control and experimental assays.

After each assay, individual copepods were photographed using either a handheld digital microscope (Amscope) or a digital camera attached to a microscope (OMAX), and their body length was measured from the cephalasome to the caudal ramus using the segmented line tool in Fiji (Schindelin et al., 2009). Since copepods were not tracked individually during assays, only the survival at the final time measurement could be used in generalized linear mixed models in order to include the length measurement for each individual as a fixed effect.

### Statistical analyses

An ANOVA was conducted for both assays to verify differential survival between treatment and control groups. To test for correlation between hypoxia tolerance and low pH tolerance, a Pearson correlation test was performed on the population-level response. Mean survival of each population was used because hypoxia and acid tolerances were not tested on the same individuals.

Linear regression was used to measure the effect size of each predictor in R version 3.6.2. Tolerance to hypoxia and low pH were measured as survival after treatment and analyzed using a generalized linear mixed effect model with a logit link for binomial distribution from the lme4 package (Bates, Mächler, Bolker, & Walker, 2015). Body length was log transformed, and all continuous variables (latitude, collection year, and log-length) were centered and scaled prior to analysis by dividing each value by twice the standard deviation (Grueber, Nakagawa, Laws, & Jamieson, 2011). Collinearity amongst predictors was checked using the car package (Fox & Weisberg, 2019) and predictors with a variance inflation factor >5 were removed from the full model. The full model for both hypoxia and acid tolerance included latitude, collection year, sex and log-length as fixed effects, and the random effects included position and group (individuals in an arena) nested within batch (arenas in the same assay). The greater sample size in the hypoxia tolerance studies provided sufficient power to allow the inclusion of all two-way interactions. After verifying and correcting for the model assumptions, the full models were dredged using the MuMIn package (Bartoń, 2019). The model with the lowest AICc was selected as the estimated best fit model, and models that had some support (ΔAICc < 5) were averaged with the estimated best fit model. Coefficient values and confidence intervals were calculated from the averaged model. To further examine the support of each predictor in the final averaged model, the weighted importance (I) of each variable—the sum of model weights across all models that included the variable—was calculated.

## Results

We measured hypoxia tolerance of 696 copepods from 6 genetically differentiated populations of the copepod *T. californicus* by measuring survival after 20 hours of hypoxia exposure and 10 hours of recovery. The hypoxia assay resulted in significantly lower survival (F_1, 694_ = 323.7, P < 0.001) than the control. Similarly, we measured acid tolerance of 472 copepods across this same set of populations by reducing the pH to 4.8 and measuring recovery after 24 hours. The low pH assay also resulted in significantly lower survival (F_1, 470_ = 99.53, P < 0.001). The Pearson correlation test for hypoxia and acid tolerance of each population suggests no correlation between the traits (t = −0.38822, P = 0.7176; Figure 1).

**Figure 1.**
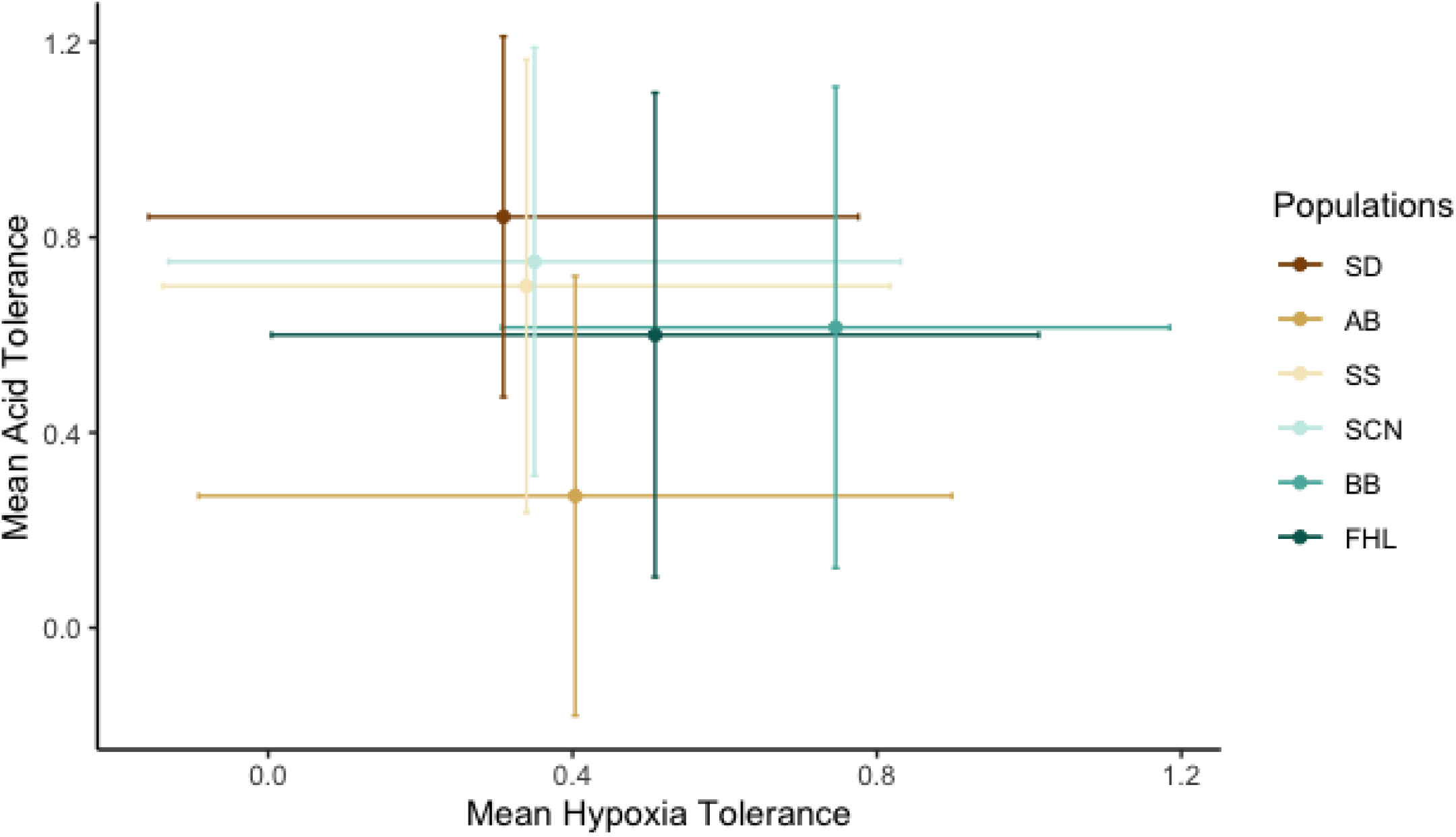
Correlation of hypoxia and pH tolerance for populations of *T. californicus*. Points are the means of the predicted values from the averaged models of hypoxia tolerance and acid tolerance by population. They are displayed with SD bars.

For hypoxia tolerance, the estimated best fit model included latitude, length and year as well as a latitude-by-year interaction, and these predictors were also the most important across all models tested (I = 1.00; Table 2). Sex was also a relatively important predictor (I ≥ 0.50), although it was not significant in the final averaged model. Tolerance to hypoxia increased with increasing latitude (Figure 2), and females were more tolerant than males (Figure 3). Tolerance to hypoxia decreased with increased length and more recent time since collection, although the latter was opposed by the effect of latitude. Older populations also tended to be northern populations, resulting in a significant latitude-by-year interaction. The remaining interactive terms were of less importance.

**Figure 2.**
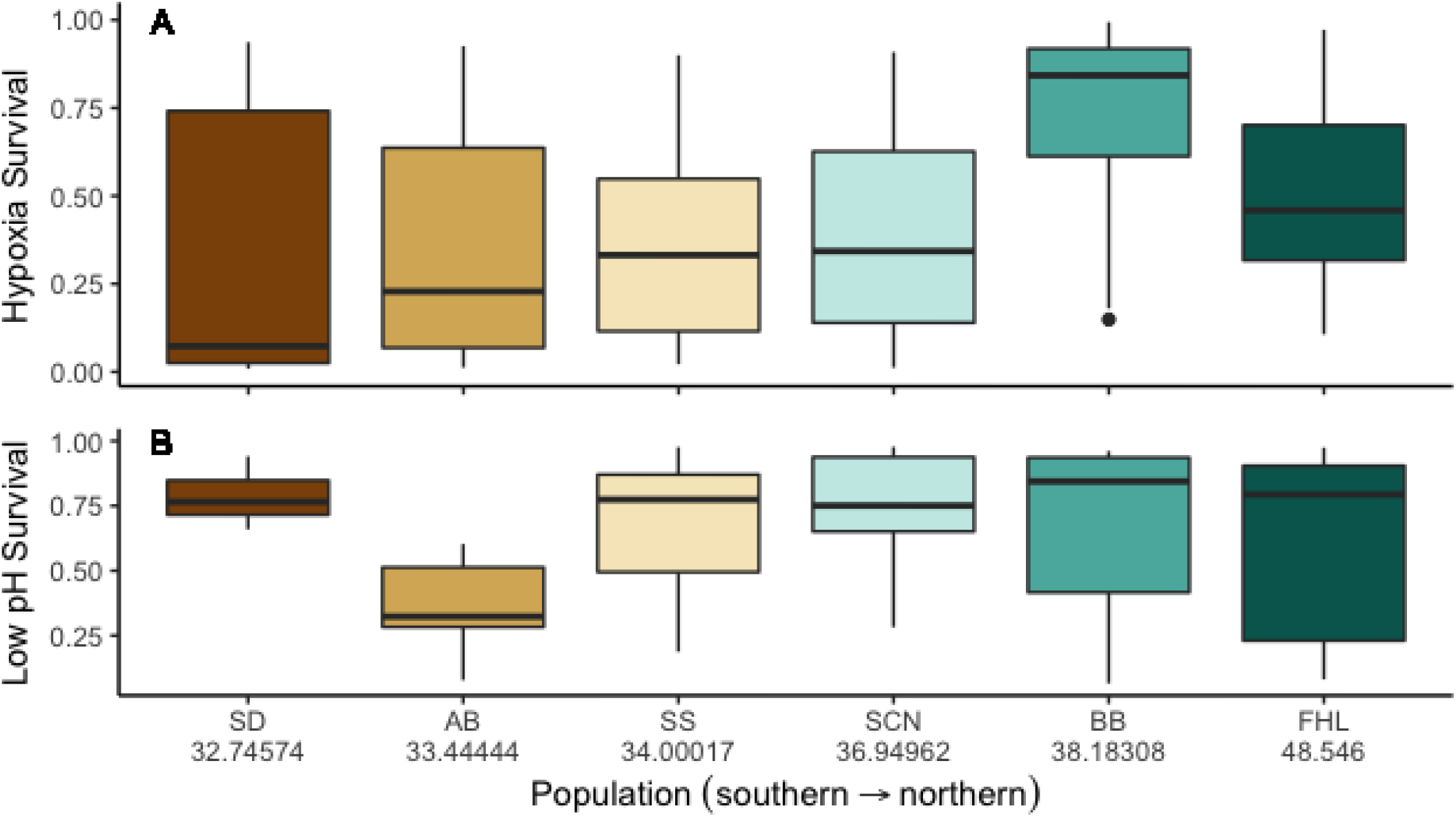
Probability of survival by population of *T. californicus* for hypoxia and pH stresses. Latitudes are written below population abbreviations in (B). Populations are ordered but not scaled by latitude. The medians are presented with the 25^th^ and 75^th^ percentiles as the box and 10^th^ and 90^th^ percentiles as whiskers.

**Figure 3.**
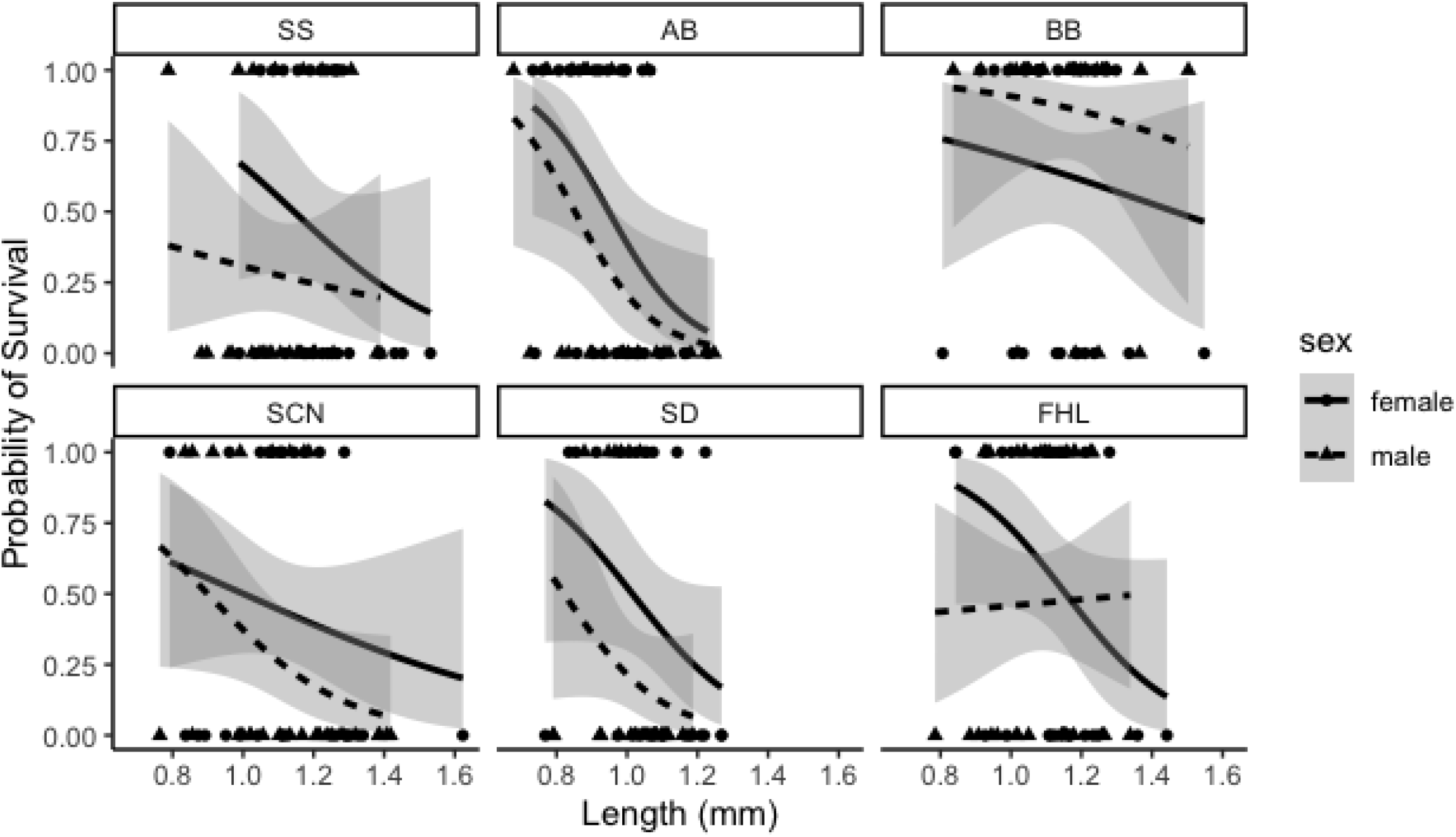
Effect of length, sex and population organized by collection date on hypoxia tolerance in *T. californicus*. The lines represent the averaged hypoxia model fit with shaded standard error bars. Points represent actual scores (1 if survived and 0 if dead). Females are solid lines and circles, and males are dashed lines and triangles. Panels are arranged from oldest collection (SS) to most recent collection (FHL). Latitude, length, collection year and latitude *×* year were significant and important effects. Sex was important but not significant.

**Table 2.** Averaged Model Values and Importance for Hypoxia Models. Significant *p*-values are written in bold. Importance is the weighted contribution of a predictor across all models compared.

| Hypoxia Model<br>Predictor<br>df = 14 | Effect size (95% Confidence Interval) | *P* | Importance |
| --- | --- | --- | --- |
| Latitude | 7.82226 (4.7412456, 10.9032754) | **<0.001** | 1.00 |
| Sex (male) | -0.2156621 (-1.1707121, 0.4057446) | 0.545342 | 0.63 |
| Length | -1.435299 (-2.1950677, -0.6755311) | **<0.001** | 1.00 |
| Collection Year | -5.127622 (-7.2924520, -2.9627913) | **<0.001** | 1.00 |
| Latitude × Sex | 0.1215641 (-0.2551879, 1.5355183) | 0.704402 | 0.25 |
| Latitude × Length | 0.07461517 (-0.3462279, 0.8645077) | 0.713292 | 0.33 |
| Latitude × Year | -6.259887 (-8.8205038, -3.6992696) | **<0.001** | 1.00 |
| Sex × Length | 0.0481855 (-0.5793483, 1.3333950) | 0.822760 | 0.20 |
| Sex × Year | -0.01750043 (-1.2015855, 0.7984095) | 0.913274 | 0.17 |
| Length × Year | 0.01313413 (-0.6190298, 0.7408713) | 0.935785 | 0.26 |

For acid tolerance, the estimated best fit model solely included sex, and sex was the only predictor with high importance across all models (Table 3). Time since collection was marginally important (0.45 < I < 0.5), and the predictors body length and latitude were less important. Males were the more sensitive sex (Figure 4), and acid tolerance increased with more recent collections. The effect for body length was indistinguishable from zero.

**Figure 4.**
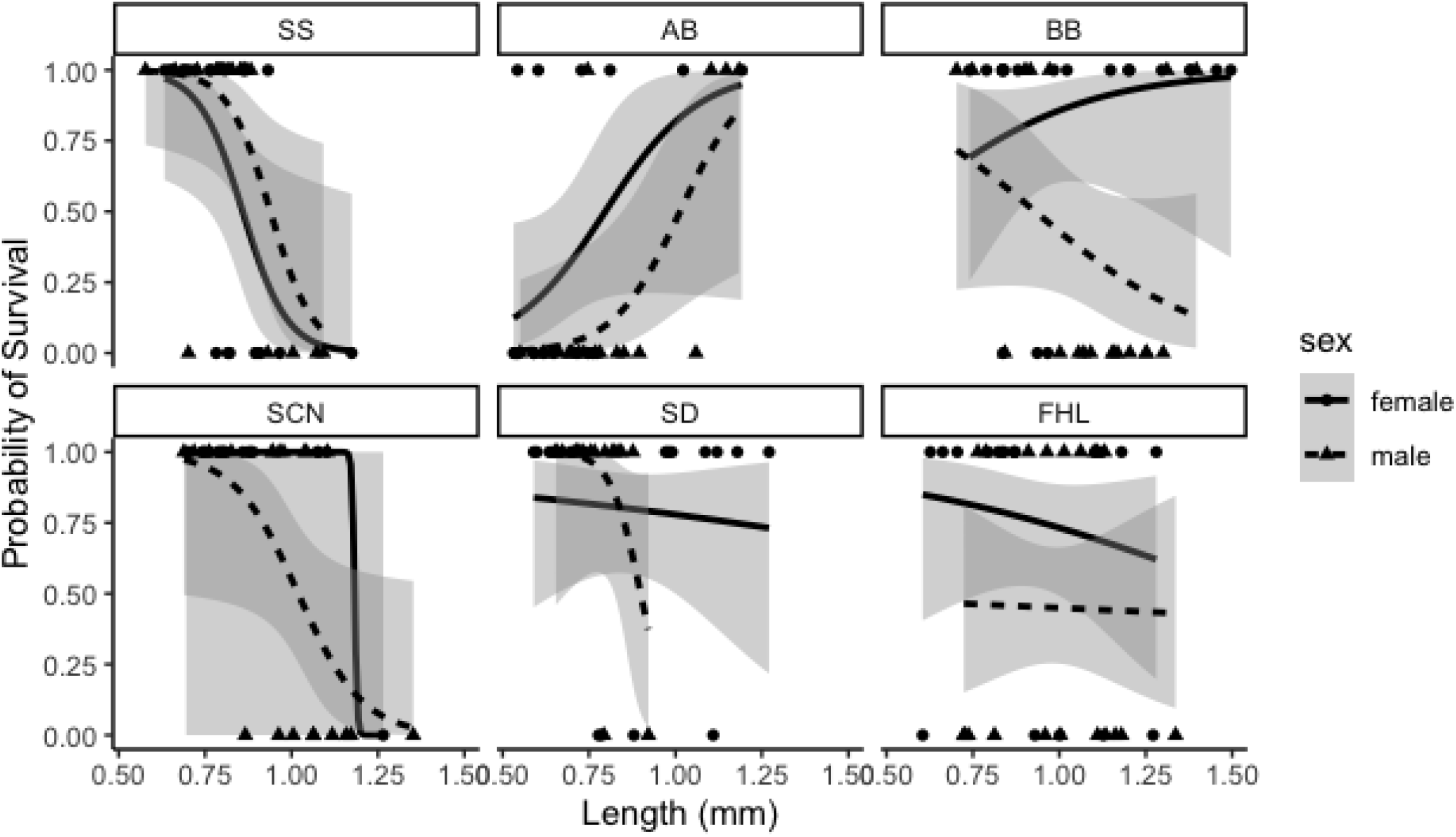
Effect of length, sex and population organized by collection date on pH tolerance in *T. californicus*. The lines represent the averaged acid model fit with shaded standard error bars. Points represent actual scores (1 if survived and 0 if dead). Females are solid lines and circles, and males are dashed lines and triangles. Panels are arranged from oldest collection (SS) to most recent collection (FHL). Sex was the only significant and important effect.

**Table 3.** Averaged Model Values and Importance for pH Models. Significant *p*-values are written in bold. Importance is the weighted contribution of a predictor across all models compared.

| pH Model Predictor<br>df = 8 | Effect size (95% Confidence Interval) | *P* | Importance |
| --- | --- | --- | --- |
| Latitude | -0.01100642 (-1.5776962, 1.4906282) | 0.9777 | 0.27 |
| Sex (male) | -1.614681 (-3.0896657, -0.3270771) | **0.0404** | 0.90 |
| Length | 0.008333252 (-1.1237287, 1.1921511) | 0.9772 | 0.26 |
| Collection Year | 0.4731268 (-0.3564342, 2.2824813) | 0.4828 | 0.49 |

## Discussion

After measuring survival in hypoxia and low pH for *T. californicus* copepods, we found no evidence of correlation in tolerance for these two stressors. This finding was surprising given how frequently these stressors covary in aquatic habitats (Hutchinson, 1957). The absence of phenotypic correlation strongly suggests that the molecular mechanisms leading to tolerance to these two traits are unrelated as well. Hypoxia tolerance in *T. californicus* likely relies on adjusting the permeability of the chitin exoskeleton to oxygen and glycolytic energy production since the species lacks respiratory pigments and the HIF-1 pathway (Graham & Barreto, 2019). In contrast, exposure to slightly acidic conditions for 24 hours resulted in a different transcriptomic response—the upregulation of oxidative stress proteins—in *Tigriopus japonicus*, a sister species of *T. californicus* (Y. H. Lee et al., 2019). It would seem that temporal covariance does not result in correlated adaptation as strongly as geographic correlation has for some other traits and species (Egea-Serrano, Hangartner, Laurila, & Räsänen, 2014; Everatt, Convey, Bale, Worland, & Hayward, 2015; Minias & Janiszewski, 2020; Pallarés, Botella-Cruz, Arribas, Millán, & Velasco, 2017), but to our knowledge the correlation between hypoxia and pH tolerance has not been directly explored previously.

The population-specific differences in hypoxia tolerance, but not acid tolerance, strongly correlated with latitude. This suggests *T. californicus* exhibit a phenotypic gradient for hypoxia tolerance, as has been documented for other environmental parameters in this species (Leong et al., 2018; Pereira et al., 2017), and any phenotypic cline in acid tolerance is significantly weaker. The latitudinal cline in thermal tolerance and hypoxia tolerance inversely covary, which was unexpected since warmer water holds less dissolved oxygen and thermal tolerance is frequently associated with hypoxia tolerance in ectotherms (Verberk & Calosi, 2012; Anttila *et al*., 2013; Schulte, 2014; McBryan *et al*., 2016; Verberk *et al*., 2018, although see also Klok *et al*., 2004; Ern *et al*., 2015; Kim *et al*., 2017). Even though coastal waters have only slight changes in dissolved oxygen (Matear & Hirst, 2003) and pH (Jiang, Carter, Feely, Lauvset, & Olsen, 2019) with latitude, the nearby water column is not representative of rock pool conditions (see Figure S3 for pH variation at one *T. californicus* location). While it is tempting to speculate about the evolutionary forces acting to produce this unexpected finding which could include, local adaptation, selection on a correlated trait, and drift, without commensurate data of latitudinal variation in oxygen and pH conditions in the high intertidal, a more definitive conclusion cannot be reached.

No other predictor changed the odds of survival for hypoxia tolerance as strongly as latitude, but sex was also an important predictor for hypoxia tolerance and the only significant predictor of acid tolerance. In both cases, males were more sensitive than females, which is consistent with previous studies in this system (Foley et al., 2019; Morgan W. Kelly, DeBiasse, Villela, Roberts, & Cecola, 2016). *T. californicus* copepods lack heterogametic sex chromosomes meaning that any sex-specific differences in tolerance cannot be explained by asymmetric inheritance of sex chromosomes but does not rule out asymmetric inheritance of the mitochondrial genome (Maklakov & Lummaa, 2013) or potential downstream effects of sex-specific endocrine expression (Rolff, 2002).

Larger individuals were more sensitive to hypoxia, but length was not a significant predictor for acid tolerance. *T. californicus* rely on cuticular diffusion for gas exchange (Graham & Barreto, 2019), so larger body size would reduce the surface area available for gas exchange. Additionally, more mass would require more oxygen to sustain metabolic functions which could explain the increased sensitivity to low oxygen (Pörtner, 2002). However, given that females and northern populations were both larger yet more hypoxia tolerant, size does not appear to affect hypoxia tolerance as strongly as other predictors which is consistent with its smaller effect size.

Time since collection was only significant for models of hypoxia tolerance. Older collections were less sensitive to hypoxia which suggests that the conditions normally employed for maintaining *T. californicus* in the lab may inadvertently be selecting for hypoxia tolerance. However, each population was collected only once, confounding our interpretation of this effect with latitude. There was a significant and negative interaction between latitude and time since collection, indicating that the effect of latitude is opposed by time in the lab. Consequently, the observed trends in hypoxia tolerance may be greater than represented in this data set, but a more clear interpretation would require repeated tests on the same population with different collection times.

*T. californicus* copepods experience a highly variable environment. Although DO was not measured in the limited environmental sampling done in this study, prior studies of *Tigriopus* habitats found similar or greater changes in pH than coastal waters with correlated decreases in DO (Morris & Taylor, 1983; Powlik, 1999; Truchot & Duhamel-Jouve, 1980). This study utilized environmental conditions outside of ranges typically experienced in native *T. californicus* habitats in order to generate observable responses rather than determining physiological limits. It is not uncommon to expose organisms to environmental conditions beyond their normal experience when assessing the limits of their tolerance (Lutterschmidt & Hutchison, 1997; Speers-Roesch, Mandic, Groom, & Richards, 2013). Both hypoxia tolerance and acid tolerance are complex traits, with multiple physiological responses that must be coordinated, and the absence of phenotype covariance in them likely underscores the absence of genetic covariance in these traits.

## Data Availability Statement

Data and code will be made freely available upon acceptance.

## Acknowledgements

We would like to thank A. Parker, M.E. Moore, K. Malinski, M. Alston, J. Lee and J.G. Kingsolver for their thoughtful feedback on the early drafts of this manuscript. We would also like to thank J. Rojas, M. Tojong, L. Scotto, H. Rendulich, and R. Pursley for their assistance in running assays and culturing copepods. This research was supported in part by a grant from the National Science Foundation [IOS-1555959 to J.G. Kingsolver and C.S.W.].

## Author Contributions

A.D. conceived and carried out the experiments. C.W. supervised and contributed to the interpretation of the results. Both authors provided critical feedback and knowledge that helped to shape the research and manuscript.

---

**Table S1.** Genetic divergence of Cytochrome B among populations of *Tigriopus californicus*. Pairwise distances were computed in MEGA (ref) using the p-distance method. Sequences for AB, BB, SC, SD, and SS were downloaded from Genbank (accession numbers included). The sequence for FHL was determined from genomic sequencing. Accession number is provided.

|  | SS | SC | BB | AB | SD | FHL |
| --- | --- | --- | --- | --- | --- | --- |
| SS<br>(GQ140992.1) | -- |  |  |  |  |  |
| SC<br>(GQ140977.1) | 0.198 | -- |  |  |  |  |
| BB<br>(GQ358922.1) | 0.206 | 0.153 | -- |  |  |  |
| AB<br>(GQ140855.1) | 0.222 | 0.206 | 0.208 | -- |  |  |
| SD_<br>GQ140645.1) | 0.214 | 0.204 | 0.218 | 0.2 | -- |  |
| FHL_(TBD) | 0.22 | 0.179 | 0.151 | 0.197 | 0.22 | -- |

**Figure S1.**
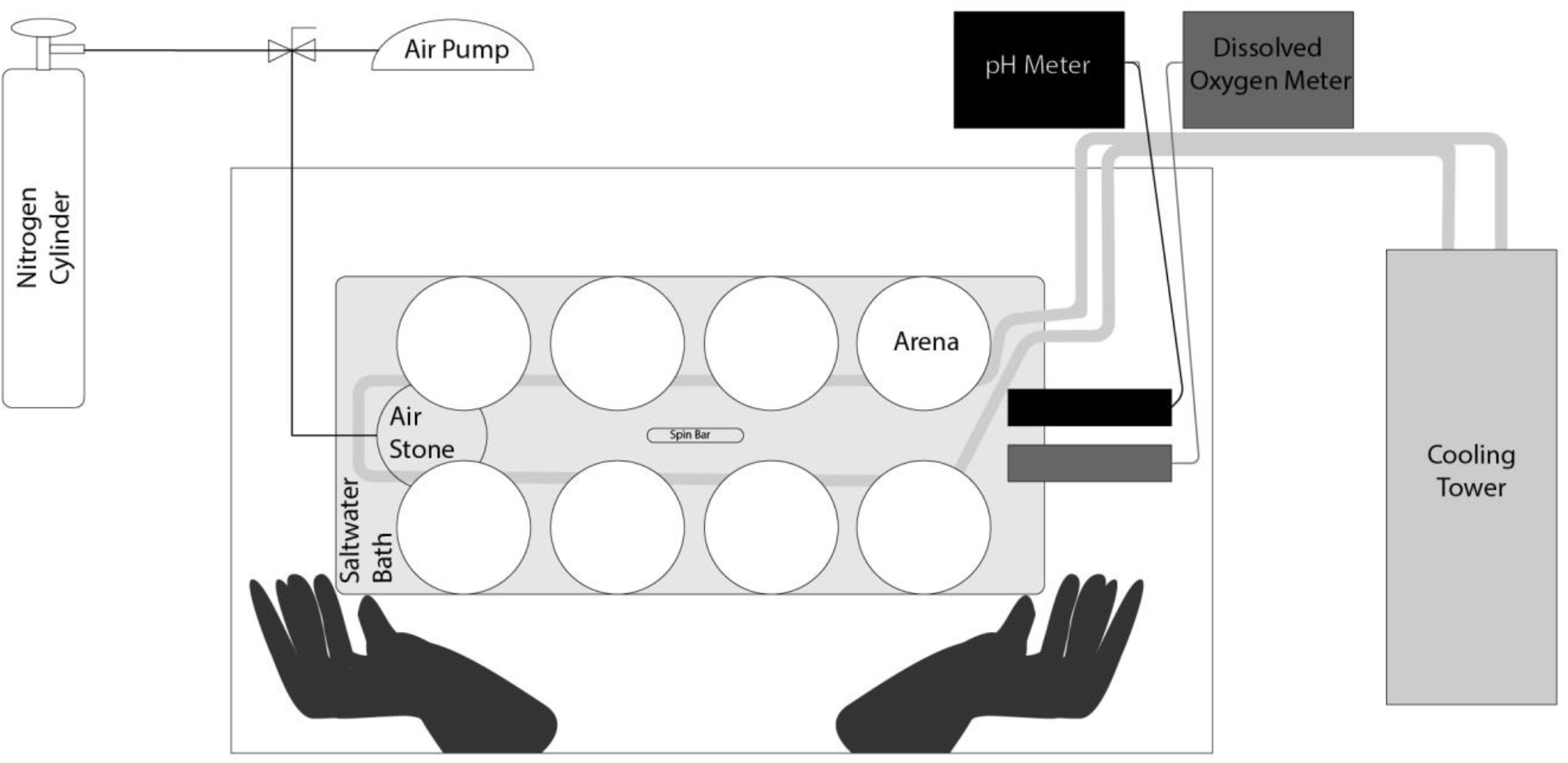
Diagram of experimental setup for hypoxia, low pH, and control assays. The glove box was constructed of ¼” plexiglass, laser cut to measure 26 in (w) x 14 in (l) x 18 in (d) with a side window held with latch clamps. Seams and cracks were sealed with epoxy.

**Figure S2.**
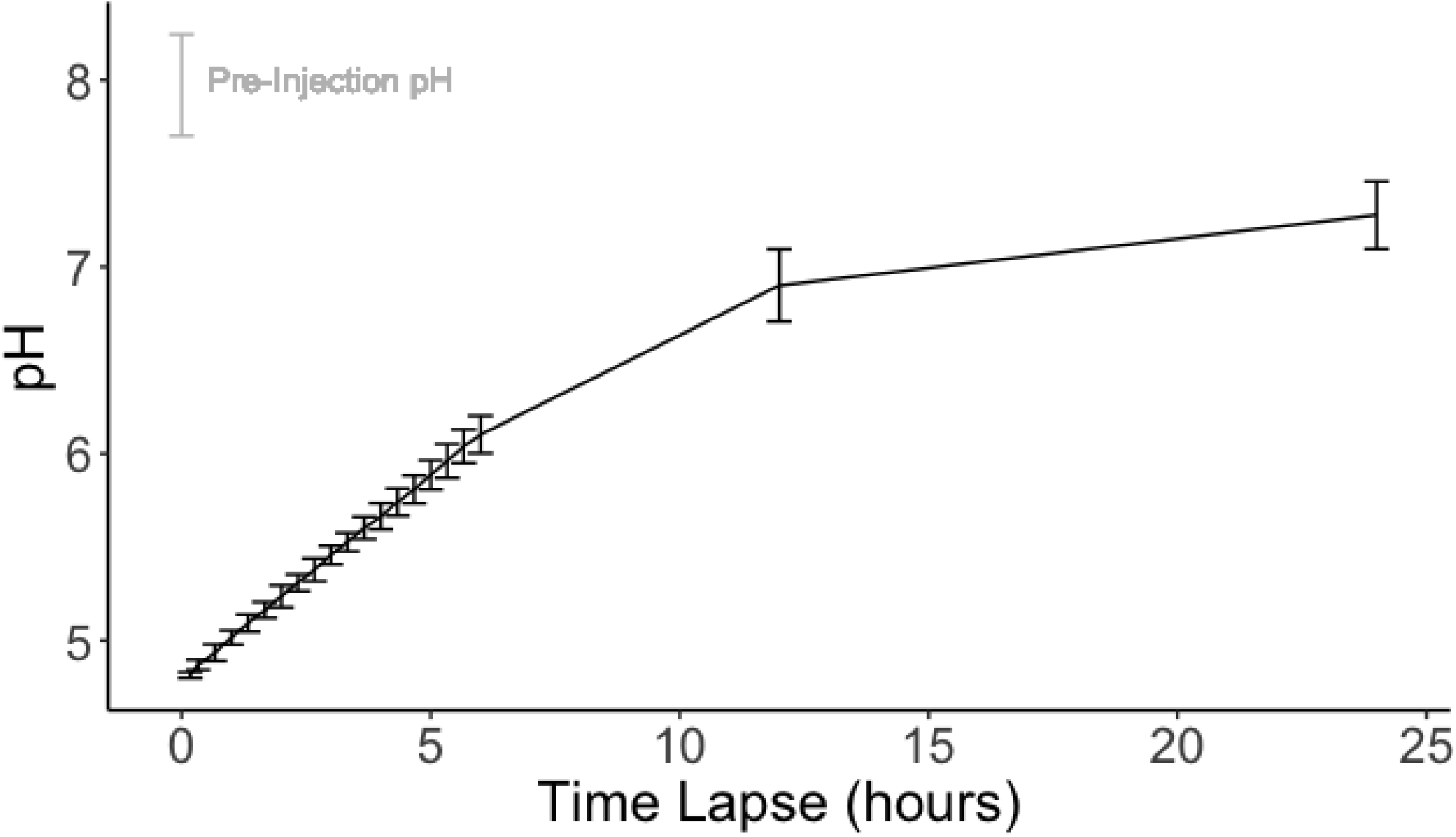
Off-Gas Rate in pH Assays. Mean pH values during low pH assays are displayed with 1 SD bars. The pH of the water bath prior to carbon dioxide injection is indicated in gray.

**Figure S3.**
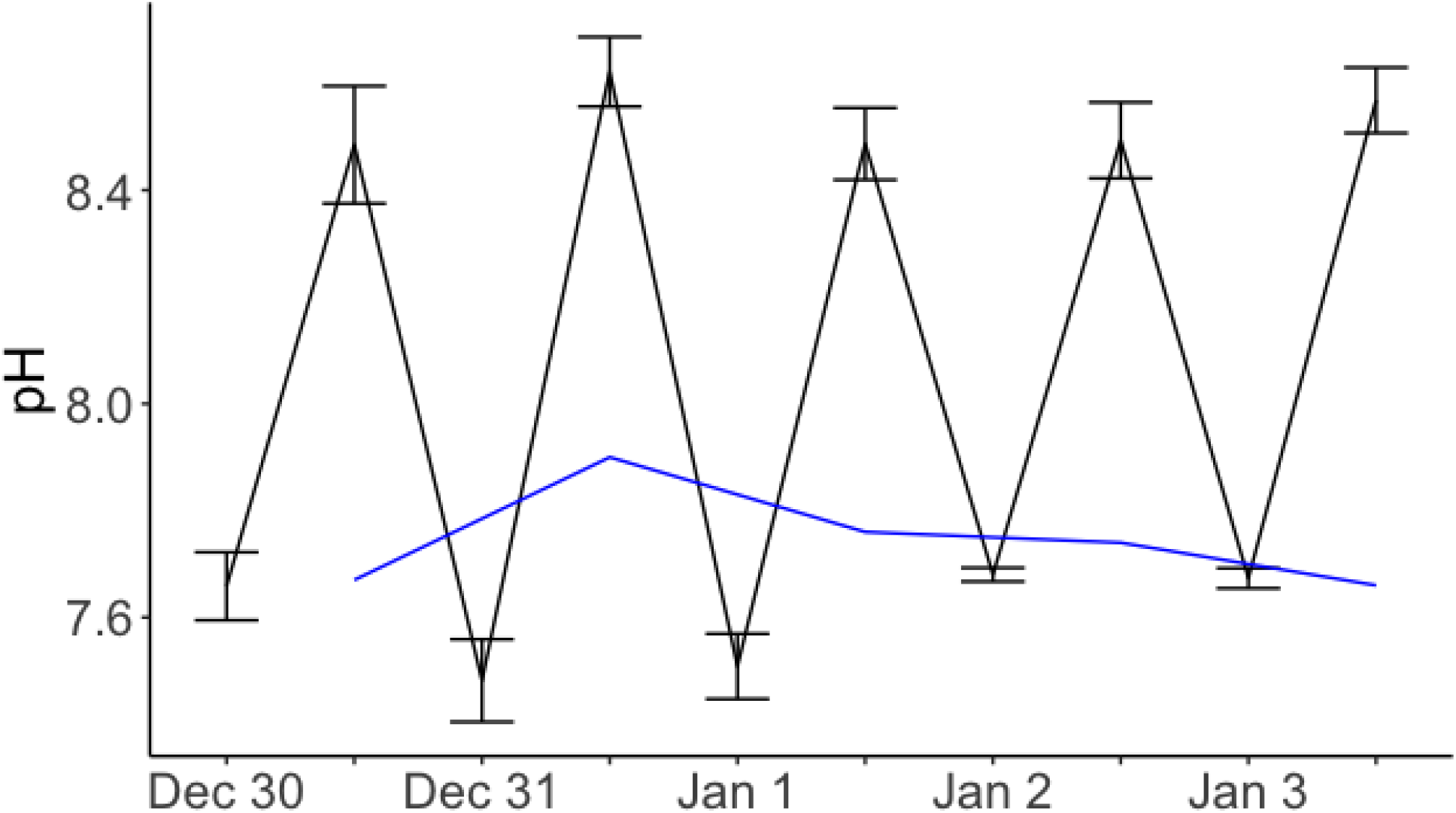
Bodega Head State Beach (BB) Rockpool and Ocean pH. All pH measurements were taken with a handheld pH meter (version). Rockpool measurements were taken at sunrise and sunset, while ocean measurements were only taken at sunset. The blackline represents rockpool values whereas the blue line represents ocean values. The values of 6 rockpools were averaged. Bars are standard errors of the mean.

## Supplementary Information

### Copepod rearing

Collections were transferred to 400 mL beakers and fed ground commercial fish flakes and natural algae growth *ad libitum*. Salinity was maintained at 35 PPT using artificial seawater, and cultures were kept in incubators at 20°C with a 12-h light/dark cycle for at least 10 generations before testing.

For hypoxia and acid assays, gravid females were removed from mass culture beakers and placed individually in 24-well plates or in groups of 10 in petri dishes and fed ground fish flakes. When nauplii were visible, the female was removed, and the nauplii were allowed to mature.

### Glovebox construction

Arenas for holding up to five copepods per trial were made from a plastic cylinder (3-inch diameter; 4-inch height) with nylon mesh (48 micron) glued to the bottom (Fig. S1). A 3-L plastic water bath connected to a cooling tower was filled with seawater and could hold 8 arenas at a time. The cooling tower maintained temperatures in the water bath at 20±1.5°C. Gases were delivered via an airstone placed at the bottom of the water bath, and a magnetic spin bar at the bottom of the water bath circulated the water among all of the arenas.

The glovebox was constructed using ¼” acrylic and laser cut following the blueprints on Dryad and sealed with clear silicone caulk. Four spring hasps were attached for the door using screws and bolts, again sealed with silicone caulk. The door was lined with self-adhesive weather stripping to insulate the door-box juncture when in use. Two 4” x 3.52” diameter coupling PVC fitting, sealed with silicone caulk, were used to secure two long kitchen gloves with hose clamps.

### Field measurements

Sunrise and sunset measurements of temperature and pH were taken for five days with an Oakton Waterproof pH Testr^®^ 30 for 6 rock pools that contained copepods at Bodega Head State Beach (38.30466, −123.06532) from Dec. 30, 2017 to Jan. 3, 2018 to provide an estimate of the spatial and diurnal variability in pH experienced by *T. californicus*. These pools were along the exposed shoreline at Bodega Head whereas collections were from Bodega Marine Lab.

From field measurements, pH was lower at sunrise (7.60±0.03) than at sunset (8.53±0.03) and differed from sunset measures of coastal water pH (7.75±0.02). Overall, pH changed an average of 0.93 units per day with a maximal change of 1.32 units (Figure S3). Diurnal changes in pH were large at nearly a 10-fold change from sunrise to sunset, much greater than what is typically experienced by marine organisms only meters away (Hofmann et al., 2011). Furthermore, these conditions frequently differed from the nearby water column, such that the timing of tidal flushing could rapidly (within minutes) alter the conditions within pools. More importantly, these findings suggest that using water column observations to estimate the physical parameters within these rock pools would be a very poor estimate of actual conditions.

